# Postzygotic isolation is stronger between sympatric species of three divergent taxa

**DOI:** 10.1101/2021.05.06.442525

**Authors:** Daniel R. Matute, Brandon S. Cooper

## Abstract

Understanding mechanisms underlying the origin of species remains a central goal of biology. Comparative studies of reproductive isolation (RI) have observed increased premating, but not postzygotic, RI between sympatric species pairs (Coyne and Orr 1989a, 1997), supporting a significant role for reinforcing natural selection in speciation. Limited data on patterns of RI have inhibited our ability to extend comparative analyses beyond model systems. Here we use an updated *Drosophila* dataset and measures of postzygotic RI from lepidopterans and toads (*Bufo*) to assess patterns of postzygotic RI in a comparative framework. Postzygotic RI is higher between sympatric species pairs of all three divergent taxa, including between some recently diverged pairs. Using the updated *Drosophila* dataset that also includes estimates of premating RI, we demonstrate that comparisons of postzygotic and premating RI depend greatly on divergence time. Both postzygotic and premating RI are strengthened between recently diverged sympatric *Drosophila* species pairs, as defined by Coyne and Orr (C&O) (Coyne and Orr 1989a) and others (i.e., *D*_Nei_ < 0.5). However, at slightly lower divergence thresholds only premating RI is significantly increased, highlighting the importance of inclusion criteria when performing comparative analyses of RI. While reinforcing natural selection against maladaptive hybridization can strengthen premating RI in sympatry, it is unlikely to contribute significantly to increases in postzygotic RI that we observe. We discuss mechanisms that may combine with reinforcement to generate increased RI in sympatry.

## INTRODUCTION

Speciation – the process in which one species splits into two – results from the accumulation of genetic differences and the evolution of reproductive isolation (RI) (Coyne and Orr 2004; Nosil 2012). In broad terms, barriers to gene flow can be classified based on when they occur in the reproductive cycle, commonly in relation to the generation of a fertilized zygote (Theodosius Dobzhansky 1937; Sobel *et al*. 2010). These barriers fall into two general categories: premating and postzygotic RI. Premating RI includes all ecological, behavioral, mechanical, and gametic incompatibilities that occur before a zygote is formed (Theodosius Dobzhansky 1937), with most focus placed on premating traits (e.g., mate discrimination). Postzygotic barriers occur after a hybrid zygote is formed and include phenotypes as extreme as hybrid sterility and inviability (Orr and Presgraves 2000; Orr 2005), but also more nuanced traits, such as hybrid behavioral defects (Delmore and Irwin 2014; Turissini *et al*. 2017; McQuillan *et al*. 2018) or delays in development (Burton 1990; Matute and Coyne 2010; Cutter and Bundus 2020).

For decades, meta-analyses have contributed to our general understanding of how RI evolves and speciation proceeds (reviewed in Matute and Cooper 2021). The earliest studies evaluated intrinsic postzygotic RI in taxa that varied in their levels of genetic differentiation (Zouros 1973; Ayala et al.1974; Wilson et al. 1974; Prager and Wilson 1975), with some studies finding no relationship between the degree of postzygotic RI and genetic distance (Zouros 1973), and others showing increased isolation for higher taxonomic groups (Ayala et al. 1974). Coyne and Orr [(Coyne and Orr 1989a, 1997); hereafter “C&O”] completed the first meta-analyses of speciation that combined estimates of genetic divergence, premating RI, and hybrid viability and fertility in *Drosophila* with phylogenetic and geographic information. Their analyses demonstrated several important results, some of which include that premating and postzygotic RI become stronger as the genetic distance between species increases and that premating RI accumulates faster in sympatry.

Analyses similar to those of C&O, and in some cases similar datasets, have revealed seminal aspects of the speciation process. Studies of the rates of evolution of RI between species pairs have enabled us to understand the role of sex chromosomes in RI (Turelli and Begun 1997; Turelli *et al*. 2014; Lima 2014), and the potential drivers of speciation (Fitzpatrick 2002; Fitzpatrick and Turelli 2006; Funk *et al*. 2006; Rabosky and Matute 2013). Expanding analyses of RI to a large variety of taxonomic groups has also confirmed that RI tends to monotonically increase with divergence in virtually all studied taxa (Gourbière and Mallet 2010; Coughlan and Matute 2020). Indeed, C&O sparked a cottage industry to study RI that has propelled some of the most important hypotheses in speciation biology (Matute and Cooper 2021).

Arguably the most influential finding of C&O was that in contrast to postzygotic RI, mate discrimination is more pronounced between sympatric than between allopatric species pairs of the same age. These results support that reinforcing natural selection against hybridization favors the evolution of premating RI when allopatric species come into secondary contact (Noor 1995; Servedio and Noor 2003; Hopkins 2013). These analyses and others support that reinforcement of premating isolation may be common during speciation (Hudson and Price 2014; Turelli *et al*. 2014), although patterns consistent with reinforcement are not always observed (Howard and Harrison 1993; Servedio and Noor 2003; Hopkins 2013). Importantly, reinforcement is expected to occur primarily at the level of premating RI (reviewed in (Servedio and Noor 2003; Coyne and Orr 2004) *cf*. (Coyne 1974 p. 19)). The availability of experimental crossing data – and specifically measurements of mate discrimination – continue to limit our ability to extend the results of C&O on reinforcement to other taxa in a comparative framework. However, studies of postzygotic RI in additional taxa do provide the opportunity to generalize a subset of the questions that C&O asked for *Drosophila*.

Here, we leverage estimates of postzygotic RI and genetic data from comparative studies of divergent *Drosophila*, lepidopteran, and toad (*Bufo*) species pairs to evaluate rates of postzygotic RI evolution between sympatric and allopatric species pairs. We observe increased postzygotic RI in sympatry for both lepidopterans and toads, including between some recently diverged species pairs. Using updated data and statistical analyses, we also find increased postzygotic RI between sympatric *Drosophila* species pairs, although this pattern emerges later in divergence than does increased premating RI. Our results support the potential for increased postmating RI to evolve in sympatry as hypothesized over a century ago (Wallace 1889; Coyne 1974). We discuss processes that may contribute to these patterns.

## MATERIALS AND METHODS

### Datasets

Our goal was to study whether the rate of evolution of postzygotic RI is similar in sympatric and allopatric animal species pairs. We gathered seven datasets that included metrics of RI (Table S1). Five of these datasets included estimates of genetic distance and the extent of geographic range overlap (i.e., whether species pairs are sympatric or allopatric). Two of these studies have fewer than five species pairs in each category, leaving only three datasets that include systematically gathered metrics of postzygotic RI in sympatric and allopatric species (Table S1).

First, we used the data on over 630 *Drosophila*-species pairs from Yukilevich (Yukilevich 2012), the most extensive compilation of measurements of reproductive RI in *Drosophila*. This compendium included all of the data from C&O (Coyne and Orr 1989a, 1997), data on postzygotic RI across *Drosophila* from Bock (Bock 1984), and new data collected by Yukilevich (Yukilevich 2012). In total, the dataset includes 288 interspecific hybridizations with estimates of genetic distance. Of these, 140 species pairs were classified as sympatric and 148 as allopatric. The second dataset was postzygotic RI in lepidopterans (data from (Presgraves 2002)). The dataset includes 212 interspecific hybridizations, 68 of which have measurements of genetic distance. Of these, 52 species pairs were classified as sympatric and 16 as allopatric. The final dataset, was postzygotic RI in toads (data from (Malone and Fontenot 2008; Castillo 2017)). This dataset includes 669 interspecific hybridizations, all of which have measurements of genetic distance (calculated from a neighbor-joining tree, (Castillo 2017)). Ninety-five species pairs were classified as sympatric and 574 as allopatric. Only the *Drosophila* dataset includes estimates of premating RI. Please note that the measurement of genetic distance between toad species (derived from a neighbor-joining tree) is not equivalent to that in *Drosophila* and Lepidopterans (Nei’s *D*), so we refrain from doing comparisons among groups between insects and anurans.

### Model fitting

We investigated the relationship between the strength of RI and the genetic distance between the parental species. While we focus on postzygotic RI in each of the three systems, we also assess prezygotic RI for the expanded *Drosophila* data set. First, we compared the fit of three models to the accumulation of RI with genetic distance: linear, logistic increase, and a four-parameter logistic (i.e., dose-response) curve. We fit the linear model using the *lm* (library *‘stats’*) function in R (R Core Team 2016) and the other two models using the *nlsLM* (library ‘*minpack*.*lm*’, (Elzhov *et al*. 2016)) function in R. The logistic models followed the form:

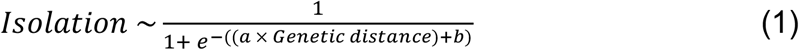

where *Isolation* is either a postzygotic or premating RI metric, *a* is the RI value when the genetic distance is zero, and *b* adjusts how quickly the probability changes with a single-unit change.

The four-parameter logistic models had the form:

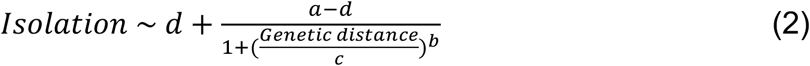

where *a* is the magnitude of isolation in a given cross at the minimum genetic distance (here where Nei’s *D* = 0), *b* is the rate of increase in RI at point *c*, the inflection point of the decay curve, and *d* is the maximum RI that can be obtained at high genetic distance. This model allows for an initial period where F1 hybrids do not suffer fitness consequences (Orr and Turelli 2001; Dagilis *et al*. 2019) and includes that RI must asymptote at a value no higher than 1 (i.e., more genetic changes contributing to isolation become redundant as two lineages cannot be more isolated than completely isolated). Since nonlinear logistic regression has difficulties optimizing the values for each of the four constants in the equations, we tried 10 starting values per constant and found the model with the lowest Akaike Information Criterion (AIC, (Akaike 1974)) with the function ‘*AIC’*, (library ‘*stats’*; (R Core Team 2016)).

To determine which of the four models best describes the relationship between isolation and genetic divergence we used AICs (Akaike 1974). To find the confidence intervals of the logistic regressions we bootstrapped the datasets 1,000 times using the function *nlsBoot* (library(‘*nlstools’*, (Baty *et al*. 2015)). Finally, we compared the confidence intervals using Wilcoxon tests of the bootstrap values for each coefficient using the function *wilcox*.*test* (library ‘*stats’*, (R Core Team 2016)).

For the subset of *Drosophila* species that had measurements of RI and Ks (genome-wide genetic distance), we similar fitted four-parameter logistic models.

### Phylogenetic independence

Ideally, our estimates of RI would be phylogenetically independent from all others. However, our *Drosophila* dataset contains multiple species pairs with phylogenetic relationships that are not evolutionarily, and thus might not be statistically, independent. To account for this lack of independence, we used clade-level and species-level sampling schemes for *Drosophila*, and a species-level sampling scheme for *Bufo* frogs (Moyle *et al*. 2004; Turissini *et al*. 2017). For the species group-level sampling scheme in *Drosophila*, we sampled a single cross from within each monophyletic clade of *Drosophila* (*affinis, ananassae, athabasca, buzzatii, melanogaster, mesophragmatica, montium, mulleri, nasuta, obscura, planitibia, pseudobscura, repleta, takahashii, virilis*, and *willistoni*; Suvorov et al. 2020) resulting in two datasets: one allopatric and one sympatric, each composed of about 16 species pairs. We subsampled the whole dataset 1,000 times and recalculated the value of *c*, the inflection point, for each iteration as described above (See Model fitting). We recorded the number of iterations that did not converge (i.e., the inferred *c* value was outside of bounds of the range of the function, in this case larger than the maximum Nei’s *D* value), but restricted comparisons between *c*_Sympatric_ and *c*_Allopatric_ to regressions that converged. We followed a similar approach for lepidopterans, subsampling by genus (*Anartia, Anthocharis, Callosamia, Choristoneura, Colias, Erebia, Heliconius, Helicoverpa, Heliothis, Hyalophora, Papilio, Phyciodes, Pieris, Pontia*, and *Ypomeneuta*). We did not use this approach for *Bufo* toads because a large proportion of the hybridizations involved species from different *Bufo sensu lato* genera (*Bufo sensu stricto, Sclerophrys, Schismaderma, Rhinella, Incillius*, and *Anaryxus*).

We also formally corrected the data using the phylogenetic mixed model approach proposed by Castillo (Castillo 2017) for the three different taxa. We fitted a generalized linear mixed model using Markov Chain Monte Carlo to study the relationship between RI and genetic distance for sympatric and allopatric species. We used the function *ginv* (library *MASS*, (Venables 2002)) to find the generalized inverse of the (1-genetic distance) matrix (Castillo 2017). We used the package *MCMCglmm* (Hadfield 2010) and fitted a linear model, in which the magnitude of isolation (premating or postzygotic for *Drosophila*, and postzygotic for Lepidopterans and *Bufo*) was the response variable, genetic distance was a predictor variable, geographic overlap (i.e., whether a species pair was sympatric or allopatric) was a fixed effect, and the phylogenetic variance matrix was a random effect. The model also included an interaction between transformed genetic distance and geographic overlap. We ran two independent MCMC chains. To determine if the model converged in each of the two chains, we used the function *gelman*.*diag* (library *coda*, (Plummer *et al*. 2006)). A chain was considered converged if all scale reduction factors for all variables (both fixed and random effects) were ≤1.1 for each of the two chains. We calculated the 95% confidence interval for the intercept and the slope using the function HPDinterval (library *coda*, (Plummer *et al*. 2006)).

### Genome-wide genetic distance in Drosophila

Nei’s *D* has been a historically important measurement of genetic distance in *Drosophila* comparative studies. Nonetheless, this metric was calculated based on single loci or allozymes (Coyne and Orr 1989a, 1997). To assess if our results were robust to the potential biases of using a single proxy of genetic divergence (i.e., Nei’s distance), we repeated our linear-model analyses using Ks, the number of synonymous differences per site calculated genome wide. We used 151 genomes for the *Drosophila* genus (Kim *et al*. 2021) and extracted a matrix of 2,791 BUSCO genes as described in (Suvorov *et al*. 2022). We calculated Ks using the count-based method, YN00 (Yang and Nielsen 2000), implemented in the program ‘yn00’ with the following parameters ‘icode = 0, weighting = 0, commonf3 × 4 = 0’. We used these pairwise distances to fit linear models in which the strength of RI was the response, whether a pair was sympatric or allopatric was a fixed effect, and Ks was a continuous variable. We used the interaction term between Ks and the geography of the species pair to assess differences in the increase of RI as divergence progresses. We used the function *lm* (library *‘stats’*) function in R (R Core Team 2016) to fit these models. These regressions are not phylogenetically corrected because the size of the dataset is small (57 species pairs).

### Comparisons of RI across divergence

Because analysis of RI across the full range of divergence may conflate conditions for co-occurrence of reproductively isolated lineages with the conditions that accelerate speciation, we next assessed RI for the most recently diverged species. Both premating and postzygotic RI data exist for *Drosophila*, enabling us to compare these measures of RI in the context of divergence. We first restricted our dataset to recently diverged *Drosophila* pairs as defined by C&O (*D*_Nei_ < 0.5) and used Wilcoxon tests to determine if postzygotic RI is strengthened at earlier stages of divergence. We took this same approach for both types of RI across the full range of Nei D. For *Bufo* pairs, we took the same approach using the available distance metric. Finally, we fit 4PL regressions to recently diverged *Drosophila* (*D*_Nei_ < 0.5) and *Bufo* pairs (NJ distance < 0.05) to assess the infection point of regressions for sympatric and allopatric pairs. We were unable to carry out these analyses for lepidopterans because the 4PL regressions did not converge when the dataset included only species pairs with Nei’s *D* < 0.5.

## RESULTS

### Increased RI in sympatry

RI increases as species diverge, and several efforts have reported a negative correlation between parental species divergence and both mating propensity (Coyne and Orr 1989a, 1997; Mendelson *et al*. 2004) and the fitness of resulting hybrids (e.g., (Bolnick and Near 2005; Lackey and Boughman 2017; Dagilis *et al*. 2019), reviewed in (Edmands 2002; Gourbière and Mallet 2010; Coughlan and Matute 2020; Matute and Cooper 2021)). We fit three types of models to study the evolution of postzygotic RI between diverging species. Table S2 shows the model parameters for this regressions. In all cases, the four-parameter logistic (henceforth abbreviated 4PL) models fit the data better than do linear or logistic models, for both allopatric and sympatric species pairs. The 4PL model meets the biological expectation of a waiting time for mutations that generate either hybrid incompatibility (Orr 1995; Orr and Orr 1996; Orr and Turelli 2001) or behavioral isolation (Mendelson *et al*. 2004), followed by a rapid accumulation of RI that eventually asymptotes.

We leveraged the information provided by our model fits and compared the values of the lower asymptote, the upper asymptote, and the inflection point for sympatric and allopatric species pairs. First, we compared the strength of RI between sympatric and allopatric species pairs of *Drosophila*. We leveraged the information provided by our model fits and compared the values of the lower asymptote, the upper asymptote, and the inflection point for sympatric and allopatric species pairs, for both postzygotic and premating RI (Figure 1A and B, respectively). Table S3 shows pairwise comparisons for the regression parameters for allopatric and sympatric species. To determine whether isolation evolved faster in sympatric species, we compared *c*, the inflection point of the 4PL model, and a proxy for how fast RI completes, for regressions using either sympatric or allopatric species. Surprisingly, we observed that for postzygotic RI, *c* is lower for sympatric than for allopatric pairs, indicating that postzygotic RI increases faster as divergence accrues between sympatric species (*c*_Sympatric-*Drosophila-*postzygotic_ = 0.439; *c*_Allopatric-*Drosophila-*postzygotic_ = 0.751; Wilcoxon rank sum test with continuity correction: *W* = 993,050, *P* < 1 × 10^−10^; Figure 1C). In Figure S1, we present the other regression parameters: *a, b*, and *d*. Consistent with previous studies (Coyne and Orr 1989a, 1997) premating RI between sympatric *Drosophila* species pairs also reaches *c* earlier than allopatric species pairs (*c*_Sympatric-*Drosophila-*premating_ = 0.059; *c*_Allopatric-*Drosophila-*premating_ = 0.318; Wilcoxon rank sum test with continuity correction: *W* = 978,080, *P* < 1 × 10^−10^; Figure 1D). These results suggest that both postzygotic and premating RI increase faster between sympatric than between allopatric species pairs. We find similar results for both postzygotic and premating RI in *Drosophila* after using stringent subsampling at the species-group level (Postzygotic: *c*_Sympatric-*Drosophila*_ = 0.38; *c*_Allopatric-*Drosophila*_ = 0.610, W = 8,342; *P* = 3.613 × 10^−10^; Premating: *c*_Sympatric-*Drosophila*_ = 0.141; *c*_Allopatric-*Drosophila*_ = 0.394, W = 16,917; *P* = 3.654 × 10^−11^; Figure S2).

**FIGURE 1.**
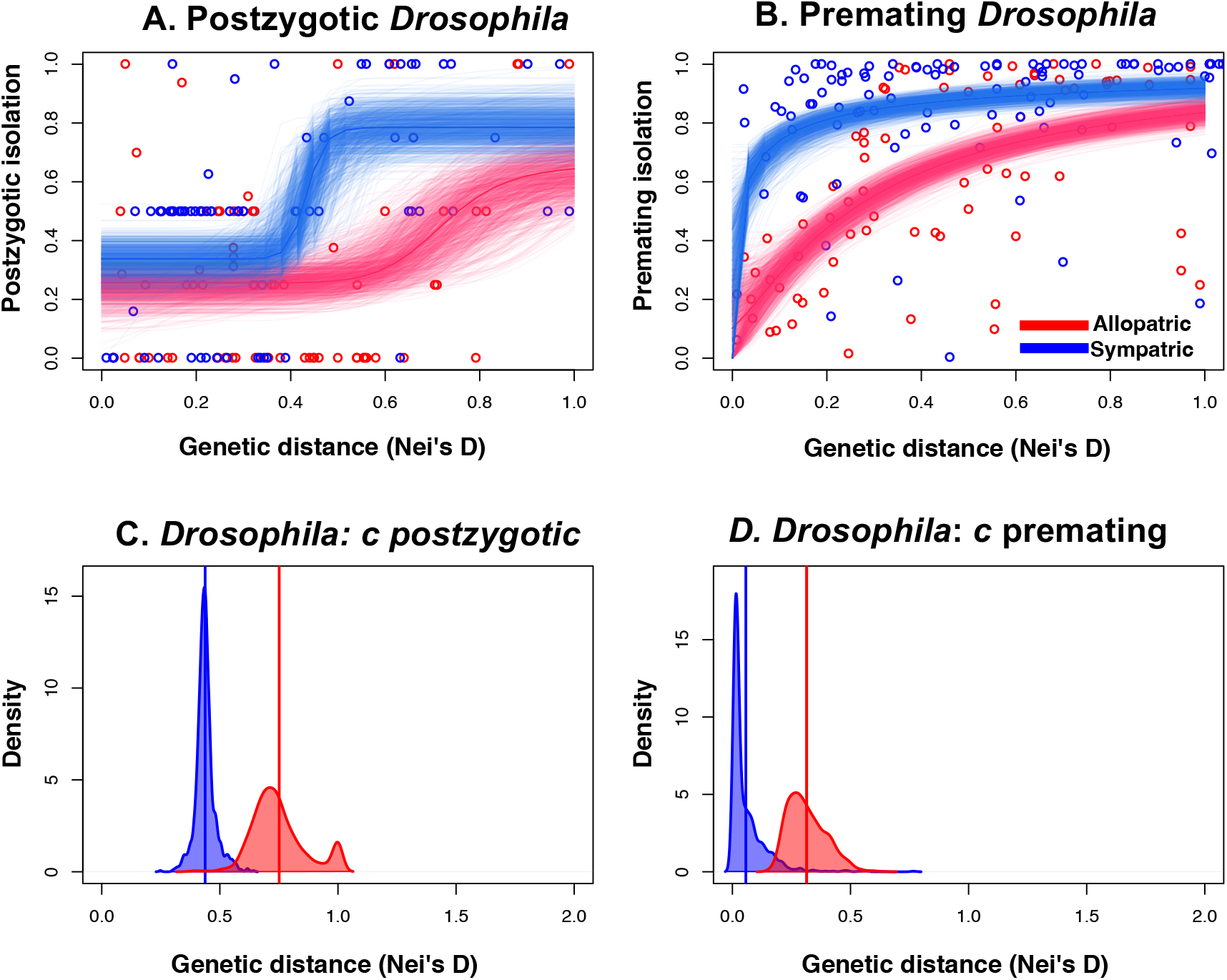
Postzygotic and premating RI evolve faster in sympatric than in allopatric *Drosophila* species pairs, but increased premating RI emerges much earlier in the process of divergence as predicted by the reinforcement hypothesis. Four-parameter logistic models for the evolution of postzygotic RI (**A**) and premating RI (**B**) between sympatric (blue) and allopatric (red) *Drosophila* species pairs. Semitransparent lines show 1,000 bootstrapped distributions. For both types of barriers, the inflection point (*c*) is at lower genetic distances for sympatric than for allopatric species pairs (premating: **C**; postzygotic: **D**). The vertical lines show the mean value of the bootstrap coefficients. Figure S1 presents the other three parameters for each of the two models.

Next, we fit a phylogenetically informed linear regression. The models included an effect for geographic origin and an interaction between origin and genetic distance. If RI evolves similarly in sympatry and allopatry, as expected for postzygotic RI under a scenario of pure-reinforcement driving the evolution of strengthened premating RI in sympatry, then effects of both origin and the interaction should be negligible. For postzygotic RI, we found that sympatric and allopatric species have a similar level of postzygotic RI (95% CI = [-0.121, 0.198], *P* = 0.660), but that postzygotic RI increases faster with genetic distance in sympatry (95%CI = [0.138, 0.699], *P* = 0.002; Table S4). For premating RI, we found that sympatric species display generally stronger premating RI than do allopatric species (95% CI=[0.245, 0.464], *P* < 0.001), but the increase of premating RI is slower in sympatric species (because of the higher intercept of sympatric species; 95% CI = [-0.323, 0.058], *P* = 0.014; Table S4). They are also consistent with the results from the uncorrected datasets, demonstrating that in *Drosophila* premating and postzygotic RI both evolve faster in sympatry.

Second, we studied the regression parameters for postzygotic RI between lepidopteran species and *Bufo* toads (Figure 2). The 4PL regression results for these taxa are similar to those in *Drosophila* (Figure 2A and B). The inflection point is lower for sympatric than for allopatric species pairs for both lepidopterans (*c*_Sympatric-Lepidopterans_ = 0.683; *c*_Allopatric-Lepidopterans_ = 1.000, *W* = 89,102, *P* < 1 × 10^−10^; Figure 2C) and *Bufo* toads (*c*_Sympatric-Toads_ = 0.028; *c*_Allopatric-Toads_ = 0.038, *W* = 130,280, *P* < 1 × 10^−10^; Figure 2D). For lepidopterans, *b* is much higher for sympatric than for allopatric species pairs (*b*_Sympatric-Lepidopterans_ = 127.60; *b*_Allopatric-Lepidopterans_ = 6.75, *W* = 130,280, *P* < 1 × 10^−10^; Figure S3), which reflects that substantial postzygotic RI is rare between allopatric species pairs even in the most divergent crosses (i.e., *Papilio xuthus* × *P. glaucus* being an exception, Nei’s *D* = 1.161). Table S3 shows pairwise comparisons for the regression parameters for allopatric and sympatric species. Subsampling by genus produces a similar result. While *c* in sympatric species is similar to the complete dataset (*c*_Sympatric-Lepidopterans_ = 0.887), *b* is lower but still positive (*b*_Sympatric-Lepidopterans_ = 5.801). Neither of these two parameters could be calculated for allopatric species because there was no increase in the magnitude of postzygotic RI over genetic distance for these species, and the 4PL regressions did not converge.

**FIGURE 2.**
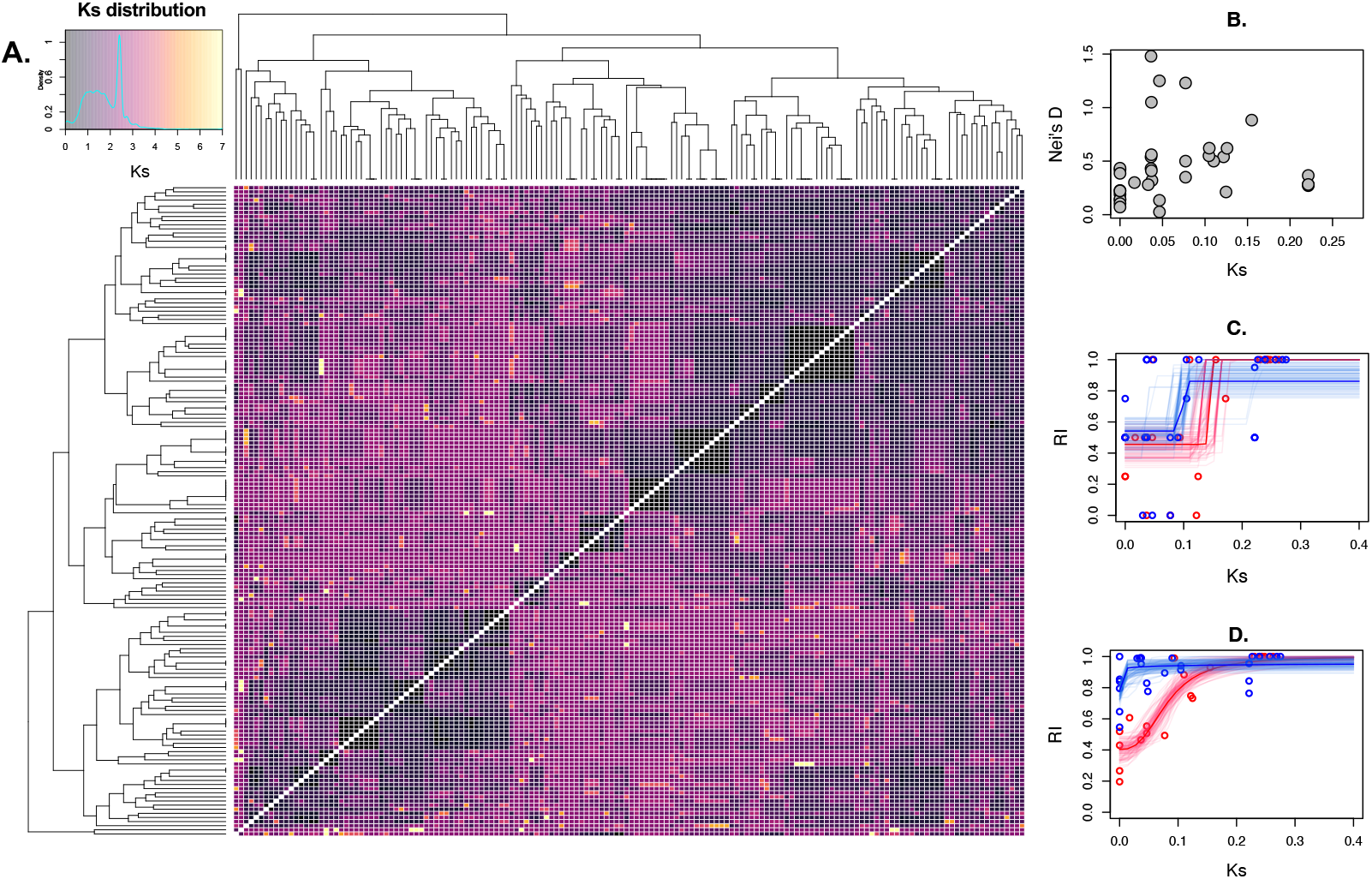
Reanalysis of the rate of accumulation of premating and postzygotic RI using Ks as a proxy of genome-wide genetic distance between species pairs. (**A**) Ks pairwise distances across the *Drosophila* genus. The inset shows the metric density across all pairs. (**B**). Ks and Nei’s D are not correlated across the entire *Drosophila* genus. Red lines show 50 allopatric bootstrap distributions for allopatric pairs, blue lines show 50 sympatric bootstrap distributions for sympatric pairs. (**C**) Faster evolution of postzygotic RI in sympatric species. (**D**) Faster evolution of premating RI in sympatric species. Color conventions are the same as in panel C.

For *Bufo, b* is higher for allopatric than for sympatric species pairs (*b*_Sympatric-Toads_ = 1.854; *b*_Allopatric-Toads_ = 3.47, *W* = 971,916 *P* < 1 × 10^−10^; Figure S3), which is consistent with the earlier inflection point for sympatric pairs and the similar asymptote of both cases (*d* = 1, Figure S3). A subsampling by subgenus was not possible for *Bufo* (See Methods).

We next carried out phylogenetically corrected linear regressions in these two groups. In the case of lepidopterans, sympatric species have slightly (non-significantly) higher postzygotic RI compared to allopatric species (95% CI = [-0.018, 0.257], *P* = 0.076). The increase of inviability with genetic distance is also faster for sympatric species than for allopatric species (95% CI = [-0.674, -0.093], *P* = 0.014; Table S5). In *Bufo*, we find that postzygotic RI is generally lower in sympatric species (95% CI = [-0.272, -0.032], *P* = 0.006), but that RI increases faster with genetic distance between sympatric species pairs (95% CI = [0.486, 3.637], *P* = 0.012; Table S5). These regressions are more limited than 4PL regressions, as linear regressions do not differentiate between the inflection point (*c*) and the rate of increase with genetic distance (*b*). Nonetheless, these results are consistent with the 4PL results in that the rate of evolution of postzygotic RI is not equivalent in allopatric and sympatric species, although additional data are required to adequately test all relevant hypotheses in a phylogenetic context in this group.

### Comparisons of RI are mostly insensitive to isolation metrics

Next, we evaluated whether the pattern of stronger premating and postzygotic RI is robust to the use of different metrics of isolation. Initial scans demonstrated that Nei’s *D*, the classic metric of genetic differentiation between *Drosophila* species used by C&O and us, is a good proxy for the extent of genome differentiation measured as synonymous differences (Ks) between species in the *melanogaster* species-group (Figure S1 in (Turissini *et al*. 2017)). To generalize this result, we calculated Ks for all pairwise combinations across the *Drosophila* genus using whole-genome data (Figure 3A). In contrast to *melanogaster* species-group comparisons, we did not find a positive correlation between Nei’s *D* and Ks across the *Drosophila* genus (Pearson’s ρ product-moment correlation = 0.087, *P* = 0.613; Figure 3B). To test whether metrics of species differentiation influence the patterns we observe, we repeated our analyses using whole-genome Ks. This necessarily reduced our dataset to 26 allopatric and 31 sympatric species pairs, but it served to fit linear models. Despite a lack of correspondence between Nei’s *D* for limited sequence data and Ks for whole genome sequence data, both postzygotic and premating RI also accumulate faster in currently sympatric species than in currently allopatric species using Ks (Figures 3C and 3D; LM Premating: interaction_*Ks*×origin_: F_1,45_ = 35.570, *P* = 3.53×10^−7^; LM Postzygotic: interaction_*Ks*×origin-_: F_1,52_= 4.943, *P* = 0.030). This indicates that the accumulation of postzygotic and premating RI we observe in *Drosophila* is not particularly sensitive to metrics of species differentiation.

**FIGURE 3.**
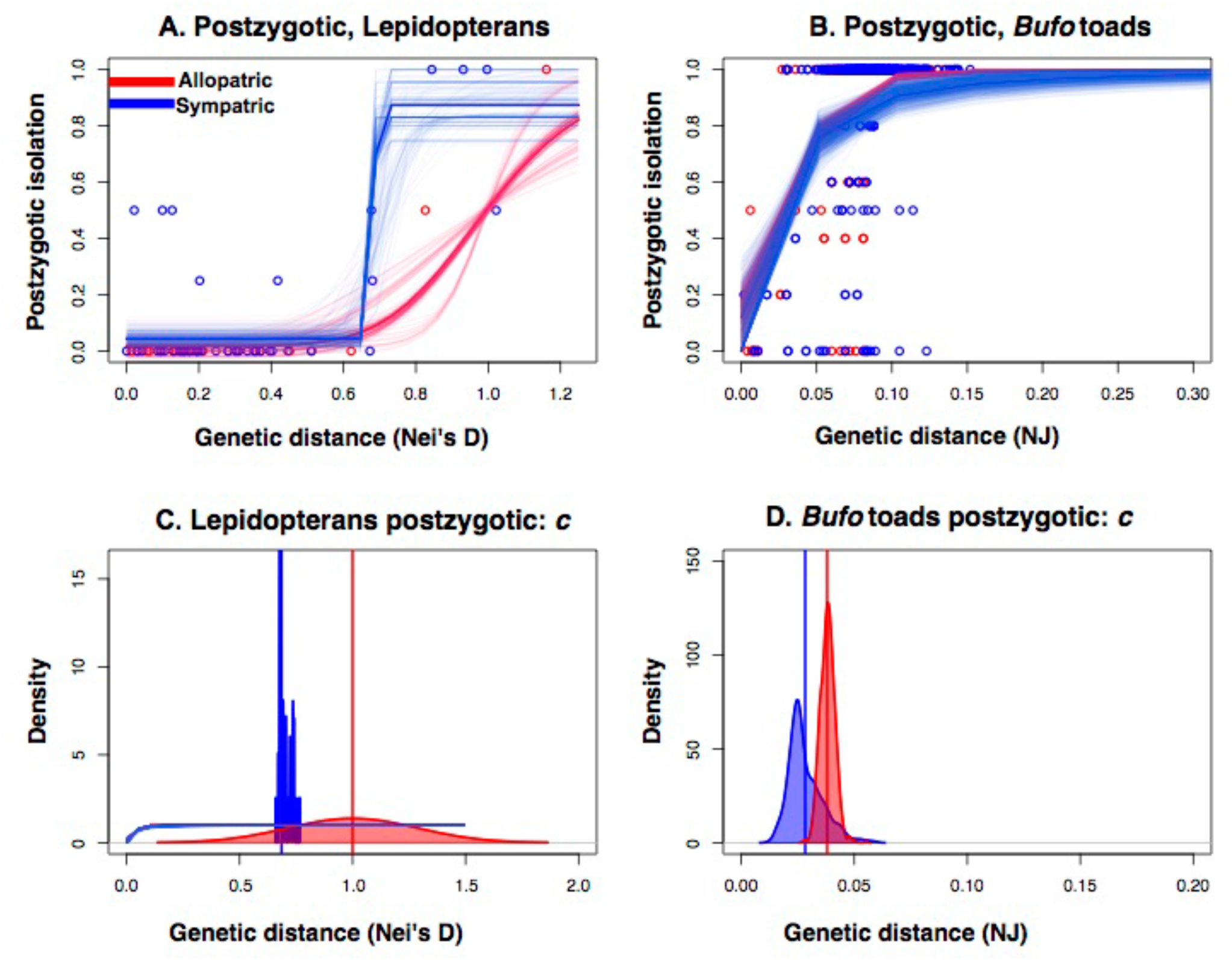
Postzygotic RI evolves faster in sympatric than in allopatric lepidopteran and *Bufo* toad species pairs. Four-parameter logistic models for the evolution of postzygotic RI between sympatric (blue) and allopatric (red) lepidopteran (**A**) and *Bufo* (**B**) species pairs. Semitransparent lines show 1,000 bootstrapped distributions. For both barriers, the inflection point (*c*) is at lower genetic distances for sympatric than for allopatric species pairs (lepidopterans: **C**; *Bufo*: **D**). The vertical lines show the mean value of the bootstrap coefficients. Figure S3 presents the other three parameters for each of the two models. Genetic distance is measured as Nei’s *D* for lepidopterans and from a neighbor-joining tree for *Bufo* (Castillo 2017).

### Comparisons of RI in sympatry depend on divergence time

Our results indicate that across the full range of divergence both postzygotic and premating RI are relatively stronger between sympatric *Drosophila* species pairs, and postzygotic RI between lepidopterans and toads tends to accumulate faster in sympatry. Because analysis of RI across the full range of divergence may conflate conditions for co-occurrence of reproductively isolated lineages with the conditions that accelerate speciation, we next assessed RI for the most recently diverged species.

We first restricted our dataset to recently diverged *Drosophila* pairs as defined by C&O (*D*_Nei_ < 0.5) to test if postzygotic RI is strengthened at earlier stages of divergence. Both postzygotic (*W* = 902.5, *P* = 0.039) and premating RI (*W* = 479.5; *P* < 0.0001) are stronger in sympatry using the C&O divergence threshold for young *Drosophila* species. However, this result is highly contingent on the arbitrary choice of how to define recently diverged species, such that with a slightly lower threshold (*D*_Nei_ = 0.47) the strength of postzygotic RI in sympatry and allopatry does not differ (*W* = 881.5, *P* = 0.05; Figure S4). In contrast, postzygotic RI between toads is significantly increased in sympatry relative to allopatry regardless of species-pair age filtering (Table S6). Fitting 4PL regressions to the most recently diverged *Drosophila* (*D*_Nei_ < 0.5) and *Bufo* pairs (NJ distance < 0.05; Figure S5) indicates that the inflection point of the regression occurs earlier for sympatric than for allopatric *Drosophila* (*c*_sympatric_ = 0.366; *c*_allopatric_ = 0.468; *W* = 812,777; *P* < 0.0001) and *Bufo* (*c*_sympatric_ = 0.029; *c*_allopatric_ > 0.5) pairs. Similar analyses were not possible for lepidopterans because the 4PL regressions did not converge when the dataset included only species pairs with Nei’s *D* < 0.5. In sum, our results indicate that comparisons of premating and postzygotic RI between sympatric *Drosophila* depend greatly on divergence time and the inclusion criteria to perform the comparative analysis.

## DISCUSSION

Research from the last three decades has revealed dozens of cases of strengthened premating RI in sympatry (reviewed in (Servedio and Noor 2003; Hopkins 2013; Hudson and Price 2014)), with evidence from experimentally evolved populations, and species, demonstrating that premating RI can evolve very rapidly if selection against hybridization is strong (Koopman 1950; Fukatami and Moriwaki 1970; Higgie *et al*. 2000; Matute 2010a; b). While comparative analyses and tests for reinforcement have emphasized increased premating RI in sympatry, postmating RI could also be strengthened in sympatry (Alfred Wallace 1889; Coyne 1974). Indeed, a handful of cases has identified instances of reinforced postmating-prezygotic RI (PMPZ) in areas of hybrid zones (Matute 2010b; Castillo and Moyle 2019). Our comparative analyses reveal patterns of increased postzygotic RI between sympatric species of three divergent taxa, suggesting that increased postzygotic RI in sympatry might be common in animals. Our results also agree with previous results that premating RI is increased between sympatric *Drosophila* species pairs, including the very most recently diverged pairs. We discuss several potential – and non-mutually exclusive – explanations for these observations.

First, stronger postzygotic RI in sympatry may result in part from species fusion and/or extinction in secondary contact (Templeton 1981a; Butlin 1987; Rhymer and Simberloff 1996). In cases when postzygotic RI is increased in sympatry, and specifically in cases when both premating and postzygotic RI are stronger in sympatry, differential fusion is a more likely driver of the pattern of enhanced RI in sympatry than reinforcement (Coyne and Orr 1989a). This idea has been disfavored because C&O did not observe elevated postzygotic RI for recently diverged sympatric *Drosophila* species, and because of the Templeton gap (i.e., the absence of young species with little reproductive isolation in sympatry; see Templeton 1981a; Coyne and Orr 1989b; Yukilevich 2012). Our observations of increased postzygotic RI in sympatry using the expanded *Drosophila* dataset and in two other animal taxa suggest these mechanisms deserve more attention in conversations about the persistence of species.

Second, while traditionally thought challenging, perhaps it is possible that postzygotic RI also evolves through pervasive reinforcing selection. This would explain the faster evolution of premating and postzygotic RI in sympatry and the strong effect of geographic overlap in both barriers. Reinforcement of postmating RI has been demonstrated in *Drosophila* (Matute 2010b; Castillo and Moyle 2019) and *Neurospora* (Turner *et al*. 2010, 2011), but in all cases the reinforced barriers act prior to zygote formation (i.e., PMPZ barriers). Even though hybrid inviability has been hypothesized to potentially evolve through selection against hybrids (Coyne 1974), reinforcement should only occur for hybrid inviability if aborting the hybrid embryos might represent an advantage to the parents. (No similar rationale has been proposed for hybrid sterility.) Since *Drosophila*, lepidopterans, and toads all show external development, there is no impetus for selection to reinforce postzygotic RI.

A third and under-discussed possibility, is that the pattern of stronger RI in sympatry is due to gene flow which might lead to a systematic underestimation of the age of sympatric species. This will disproportionately affect recently diverged sympatric species as the extent of gene flow seems to be negatively correlated with the divergence age between the parental species (Kronforst *et al*. 2013; Hamlin *et al*. 2020). This possibility might affect our inference of stronger postzygotic RI in sympatry and specifically create the appearance of stronger RI at low genetic distances. We argue this is an unlikely possibility. Assessments using phylogenetically-based methods suggest that gene exchange seems to be pervasive. The percentage of the genome that shows evidence of gene exchange among *Drosophila* species pairs ranges between 0.1% and 5% (Kulathinal *et al*. 2009; Garrigan *et al*. 2012; Turissini and Matute 2017). Nonetheless, coalescent-based methods suggest that there are no detectable differences in the proportion of introgression between currently sympatric and allopatric species (Yusuf *et al*. 2022), which in turn questions the fidelity between the current geographical and putative historical ranges. Additionally, the vast majority of introgression is purged rapidly in *Drosophila* hybrid populations (i.e., within the first ten generations) after admixture (Veller *et al*. 2019; Matute *et al*. 2019; Suvorov *et al*. 2020). The effect of introgression on point measurements of differentiation in currently allopatric and currently sympatric species pairs deserves a systematic treatment.

Our analyses suggest that a combination of processes contribute to strengthened postzygotic RI in sympatry. Considering the timescale of increased premating RI and postzygotic RI between sympatric *Drosophila* species (and several other results we present), we hypothesize that species fusion and/or extinction in secondary contact may contribute significantly to increased postzygotic RI in sympatry (Templeton 1981b). While reinforcing natural selection against maladaptive hybridization contributes to strengthened premating RI in sympatry, differential fusion and/or extinction could plausibly also contribute to this pattern, in addition to increased postzygotic RI. As noted by C&O (Coyne and Orr 1989a) “*enhanced premating RI could result from a process of fusion or extinction in sympatry*” because “*any factor that reduces gene flow should inhibit fusion or extinction*” (pg. 376). A concerted effort to compare the prevalence of strengthened premating and postzygotic RI in sympatry – and its potential causes across divergent taxa – will contribute to a deeper understanding of these patterns. We expect that those analyses will continue to reveal that no single mechanism underlies elevated RI in sympatry.

## Acknowledgements

We thank J. M. Coughlan and A. Dagilis for comments and J. M. Good for helpful discussions. Research reported in this publication was supported by the National Institute of General Medical Sciences of the National Institutes of Health (NIH) under Award Number R35GM148244 to DRM and R35GM124701 to BSC. The content is solely the responsibility of the authors and does not necessarily represent the official views of the NIH.

