## Supplementary material for "Postzygotic isolation is stronger between sympatric species of three divergent taxa": Figures S1-S6, Tables S1-S9

**This PDF file includes:**

Figures S1 to S6

Tables S1 to S9

**FIGURE S1.** 4PL regression parameters for *Drosophila* ……………………..…………**2**

**FIGURE S2.** *Drosophila* 4PL regression parameters after phylogenetic subsamplings at the species group level……………………………………………………………………**3**

**FIGURE S3.** 4PL regression parameters for Lepidopterans and *Bufo* frogs……….……**4**

**FIGURE S4.** Comparisons of RI depend greatly on divergence time……………………..**5**

**FIGURE S5.** Postzygotic isolation evolves faster in sympatric species than in allopatric species when the data set is restricted to 'young' species……………..……...……….....**6**

**TABLE S1.** Datasets that have compiled data on the magnitude of reproductive isolation in multiple species pairs of animals..........……......................…………………...**7**

**TABLE S2.** 4PL models fit the data better than linear regressions……………………...**8**

**TABLE S3.** Pairwise comparisons between coefficients of the four-parameter logistic model between sympatric and allopatric species…………………..…………………...**9**

**TABLE S4.** Coefficients for a phylogenetically informed linear regression for premating and postzygotic isolation between *Drosophila* species pairs……………………………..**11**

**TABLE S5.** Regression coefficients from a phylogenetically informed regression for postzygotic isolation between Lepidopteran and *Bufo* species pairs……………………**12**

**TABLE S6.** Young species show faster accumulation of postzygotic isolation when they are sympatric than when they are allopatric in *Bufo* frogs………………………………...**13**

**FIGURE S1. 4PL regression parameters for *Drosophila*.** The density plots correspond to *a* (left), *b* (middle), and *d* (right) for premating isolation (top) and postzygotic isolation (bottom). Blue: sympatric; red: allopatric. The values of *c*, the fourth parameter of the regression, are shown in Figure 1.

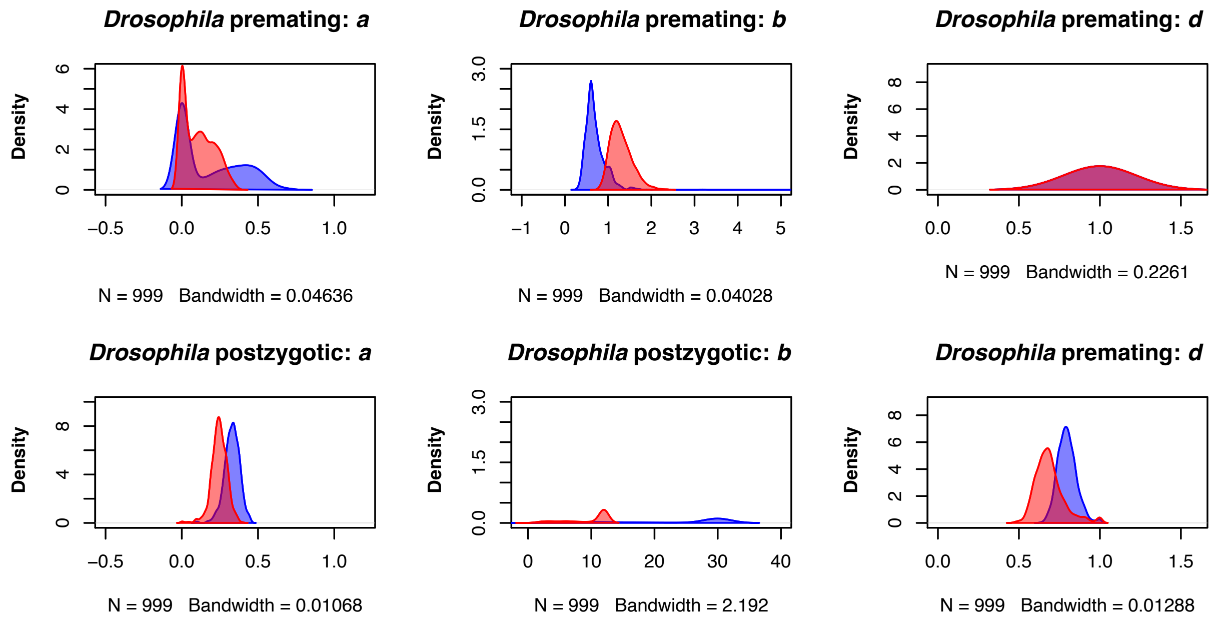

**FIGURE S2. *Drosophila* 4PL regression parameters after phylogenetic subsamplings at the species-group level.** The density plots correspond to *a* (left), *b* (middle left), *c* (middle right), and *d* (right) for premating isolation (top) and postzygotic isolation (bottom). Blue: sympatric; red: allopatric.

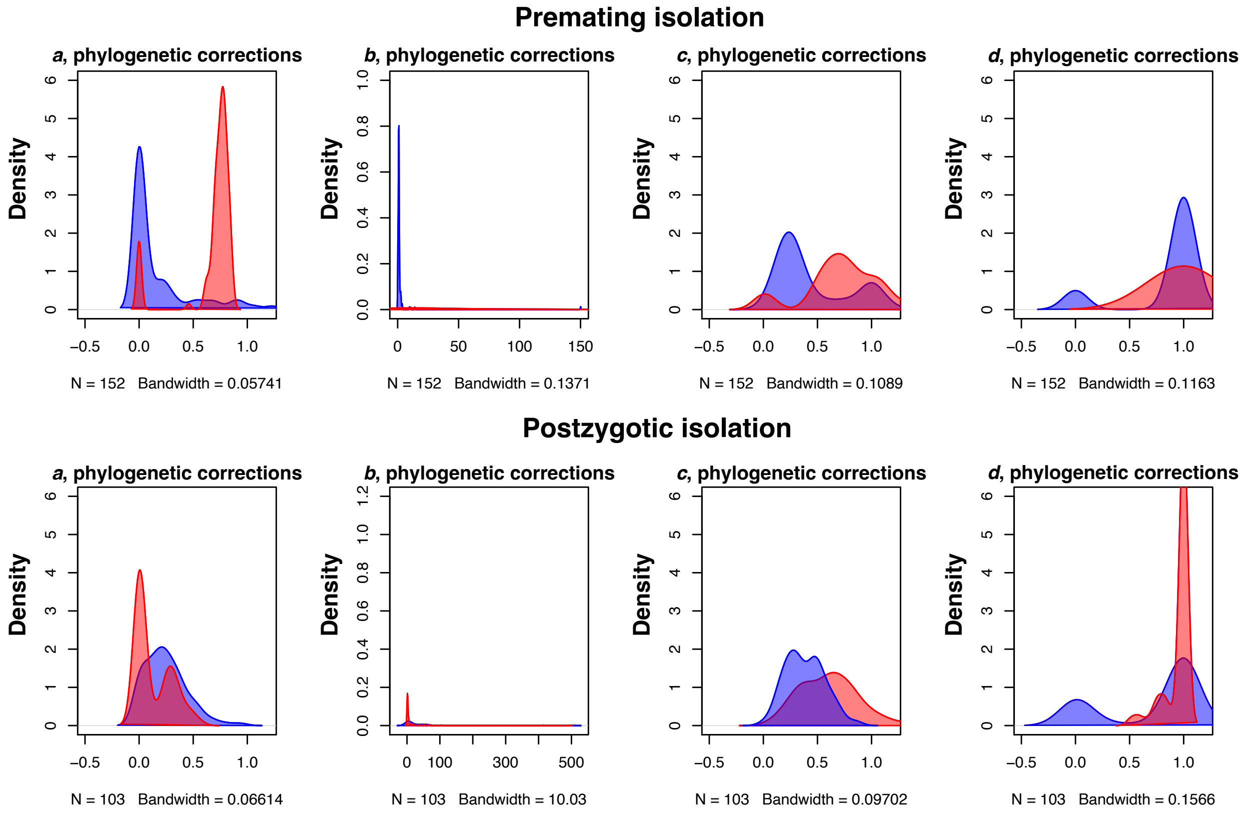

**FIGURE S3. 4PL regression parameters for Lepidopterans and *Bufo* frogs.** The density plots correspond to *a* (left), *b* (middle), and *d* (right). Blue: sympatric; red: allopatric. The values of *c*, the fourth parameter of the regression, are shown in Figure 2.

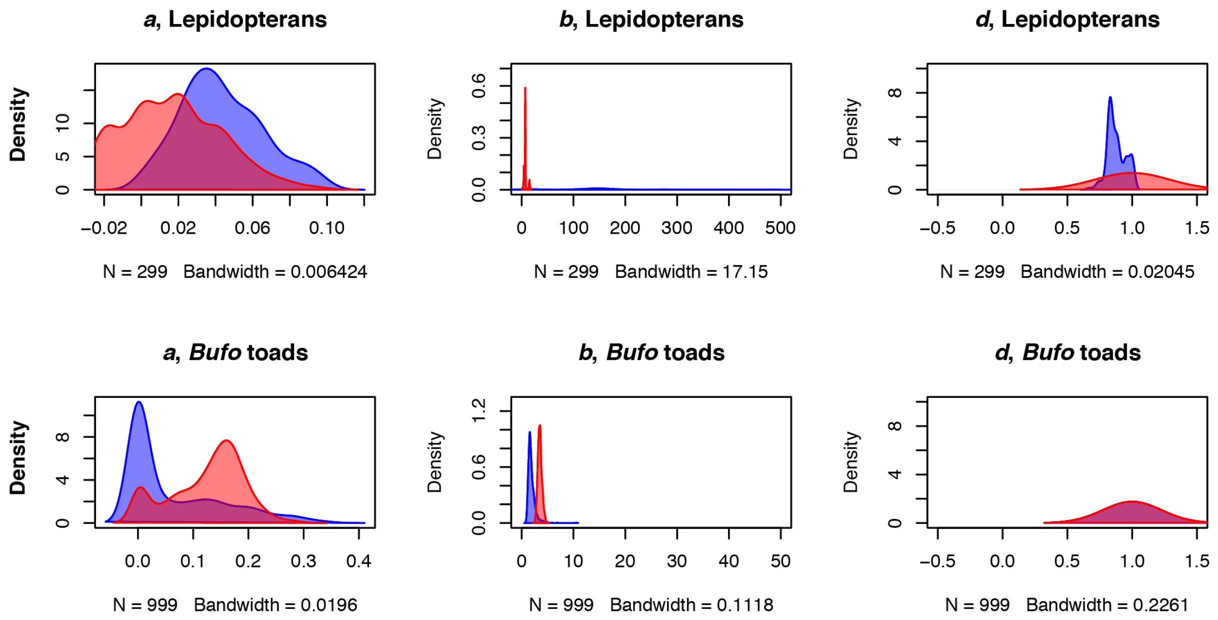

**FIGURE S4. Comparisons of RI depend greatly on divergence time.** Wilcoxon tests comparing the strength of premating (black) and postzygotic (purple) RI in sympatry and allopatry when using different thresholds to define young species. The solid horizontal line indicates the threshold of significance defined by Coyne and Orr (1, 2). Shaded intervals represent 95% confidence intervals.

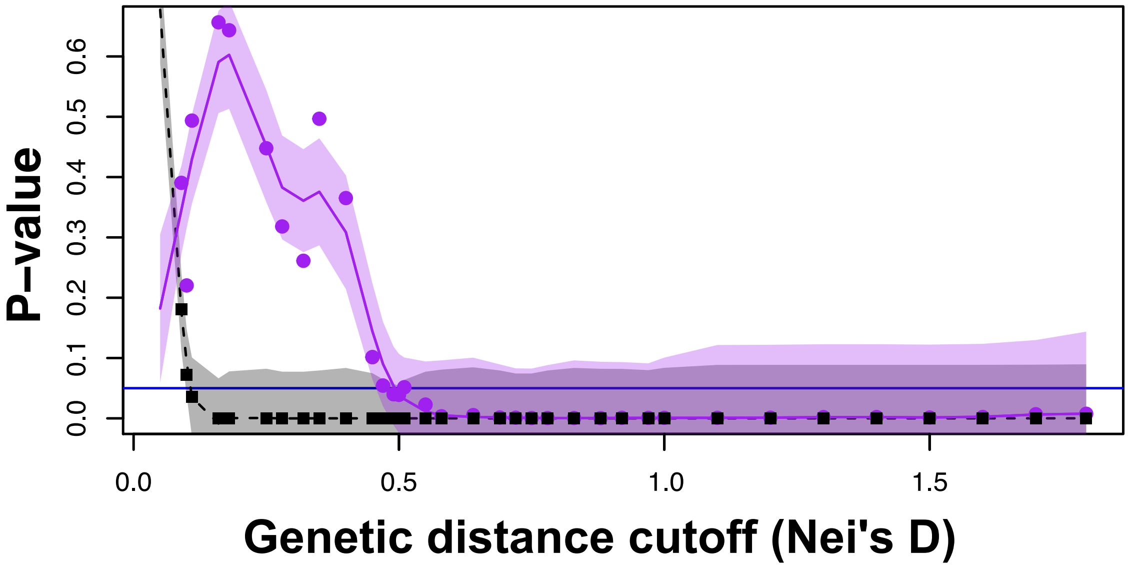

**FIGURE S5. Postzygotic isolation evolves faster in sympatric species than in allopatric species when the data set is restricted to ‘young’ species.** For these analyses we defined young as the species with the third lowest divergence time (i.e., Nei’s *D* < 0.5 for *Drosophila* and NJ distance < 0.05 for *Bufo)*. 4PL regressions in the youngest *Drosophila* (Nei’s *D* < 0.51) and youngest *Bufo* species (Nei’s *D* < 0.51) show similar results to the full range of divergence. In both cases (young species and full divergence), sympatric species pairs show an earlier inflection point. Note that we did not do the same analyses for Lepidopterans as the 4PL regressions did not converge in datasets that did not include all the data.

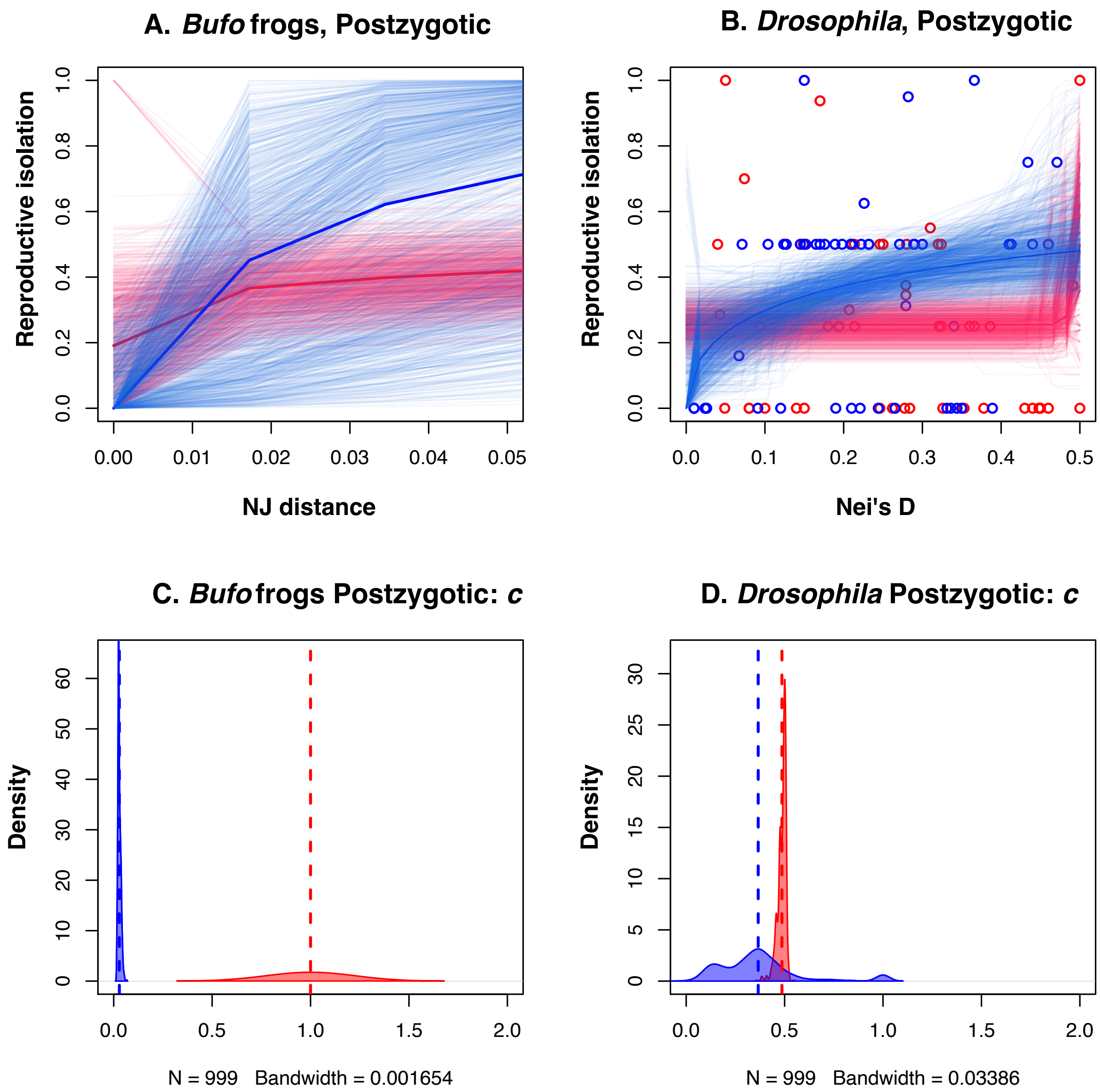

### SUPPLEMENTARY TABLES

**TABLE S1. Datasets that have compiled data on the magnitude of reproductive isolation in multiple species pairs of animals.**

| | ***N*** | **Premating isolation** | **Postzygotic isolation** | **Distance** | **Geographic status** | **Reference** |
|---|---|---|---|---|---|---|
| ***Drosophila*** | 630 | X | X | X | X | (6) |
| **Toads** | 669 | | X | X | X | (7, 8) |
| **Lepidoptera** | 212 | | X | X | X | (9) |
| ***Heliconius*** | 12 | X | X | X | | (10) |
| **Birds** | 407 | | X | X | | (11) |
| **Sticklebacks** | 5 | X | X | X | | (12) |
| **Centrarchid fish** | 130 | | X | X | | (13) |

**TABLE S2. 4PL models fit the data better than linear regressions.** Akaike information criterion for each fitted model.

| Taxon | Barrier | Linear | Logistic | 4PL |
|---|---|---|---|---|
| ***Drosophila*** | **Premating. Allopatric** | 3.111 | 7.167 | -19.557 |
| ***Drosophila*** | **Premating. Sympatric** | -44.350 | -45.059 | -53.562 |
| ***Drosophila*** | **Postzygotic. Allopatric** | 53.975 | 53.383 | 50.630 |
| ***Drosophila*** | **Postzygotic. Sympatric** | 36.043 | 138.178 | 30.949 |
| **Lepidopterans** | **Postzygotic. Allopatric** | -13.427 | -16.391 | -19.366 |
| **Lepidopterans** | **Postzygotic. Sympatric** | -11.07777 | 5.3061 | -44.391 |

**TABLE S3. Pairwise comparisons between coefficients of the four-parameter logistic model between sympatric and allopatric species.** To assess the significance of the difference between coefficients we compared bootstrapped distributions using a Wilcoxon test.

| | Coefficient | $Coef_{sympatric}$ | $Coef_{allopatric}$ | Wilcoxon test |
|---|---|---|---|---|
| **Premating, *Drosophila*** | *a* | 0.184 (0.211) | 0.115 (0.102) | *W* = 468,480; *P* = 0.015 |
| **Premating, *Drosophila*** | *b* | 0.740 (0.517) | 1.316 (0.270) | *W* = 949,860; *P* < 0.0001 |
| **Premating, *Drosophila*** | *c* | 0.059 (0.083) | 0.318 (0.085) | *W* = 978,080; P < 0.0001 |
| **Premating, *Drosophila*** | *d* | 1 (0) | 1 (0) | *W* = 499,000; *P* = NA |
| **Postzygotic, *Drosophila*** | *a* | 0.329 (0.049) | 0.242 (0.050) | *W* = 93,010; *P* < 0.0001 |
| **Postzygotic, *Drosophila*** | *b* | 22.843 (9.801) | 9.402 (3.532) | *W* = 164,870; *P* < 0.0001 |
| **Postzygotic, *Drosophila*** | *c* | 0.439 (0.044) | 0.751 (0.114) | *W* = 993,050; *P* < 0.0001 |
| **Postzygotic, *Drosophila*** | *d* | 0.799(0.061) | 0.682 (0.091) | *W* = 126,760; *P* < 0.0001 |
| **Postzygotic, Lepidopterans** | *a* | 0.015 (0.025) | 0.043 (0.021) | *W* = 17155; *P* < 0.0001 |
| **Postzygotic, Lepidopterans** | *b* | 7.252 (2.955) | 222.696 (426.678) | *W* = 1,189; *P* < 0.0001 |
| **Postzygotic, Lepidopterans** | *c* | 1 (0) | 0.686 (0.012) | *W* = 89,401; *P* < 0.0001 |
| **Postzygotic, Lepidopterans** | *d* | 1(0) | 0.872 (0.069) | *W* = 85,364; *P* < 0.0001 |
| **Postzygotic,** | *a* | 0.062 (0.087) | 0.124 (0.065) | *W* = 739,185; |

| | | | | |
|---|---|---|---|---|
| ***Bufo* toads** | | | | $P < 0.0001$ |
| **Postzygotic, *Bufo* toads** | *b* | 1.854(0.699) | 3.520 (0.414) | $W = 971{,}916$; $P < 0.0001$ |
| **Postzygotic, *Bufo* toads** | *c* | 0.028 (0.008) | 0.038 (0.003) | $W = 877{,}636$; $P < 0.0001$ |
| **Postzygotic, *Bufo* toads** | *d* | 1 (0) | 1 (0) | $W = 499{,}000$, $P$ = NA |

**TABLE S4. Coefficients for a phylogenetically informed linear regression for premating and postzygotic isolation between *Drosophila* species pairs.** The confidence intervals are based in 5 different chains.

| **Isolation** | **Coefficient** | **Isolation mean** | **95% CI** | **Effective sample size** | **pMCMC** |
|---|---|---|---|---|---|
| **Premating** | (Intercept) | 0.417 | [0.253, 0.616] | 888.760 | <0.001 |
| **Premating** | Genetic Distance | 0.297 | [0.179, 0.414] | 11.710 | <0.001 |
| **Premating** | Sympatry (Sympatric-Allopatric) | 0.364 | [0.245, 0.464] | 575.470 | <0.001 |
| **Premating** | Genetic Distance × Sympatry (Sympatric-Allopatric) | -0.184 | [-0.323, -0.058] | 1,000.00 | 0.014 |
| **Postzygotic** | Intercept | 0.214 | [-0.018. 0.446] | 910.500 | 0.084 |
| **Postzygotic** | Genetic Distance | 0.306 | [0.087, 0.530] | 704.400 | 0.004 |
| **Postzygotic** | Sympatry (Sympatric-Allopatric) | 0.034 | [-0.121, 0.198] | 1,000.00 | 0.660 |
| **Postzygotic** | Genetic Distance × Sympatry (Sympatric-Allopatric) | 0.388 | [0.138, 0.699] | 984.50 | 0.002 |

**TABLE S5. Regression coefficients from a phylogenetically informed regression for postzygotic isolation between Lepidopteran and *Bufo* species pairs.** The confidence intervals are based in 5 different chains.

| Group | | Postzygotic mean | 95% CI | Effective sample size | pMCMC |
|---|---|---|---|---|---|
| **Lepidopterans** | (Intercept) | -0.100 | [-0.392, 0. 273] | 1,000.0 | 0.488 |
| **Lepidopterans** | Genetic Distance | 0.906 | [0.670, 1.125] | 892.2 | <0.001 |
| **Lepidopterans** | Sympatry (Sympatric-Allopatric) | 0.123 | [-0.018, 0.257] | 1,000.0 | 0.076 |
| **Lepidopterans** | Genetic Distance × Sympatry (Sympatric-Allopatric) | -0.382 | [-0.674, -0.093] | 1,000.0 | 0.014 |
| ***Bufo* frogs** | (Intercept) | 0.502 | [0.294, 0.691] | 1,000.0 | <0.001 |
| ***Bufo* frogs** | Genetic Distance | 4.849 | [3.668, 5.950] | 1,000.0 | <0.001 |
| ***Bufo* frogs** | Sympatry (Sympatric-Allopatric) | -0.161 | [-0.272, -0.032] | 1,000.0 | 0.006 |
| ***Bufo* frogs** | Genetic Distance × Sympatry (Sympatric-Allopatric) | 1.938 | [0.486, 3.637] | 1,000.0 | 0.012 |

**TABLE S6. Recently diverged species show faster accumulation of postzygotic isolation when they are sympatric than when they are allopatric in *Bufo* frogs.** Wilcoxon tests comparing the strength of postzygotic isolation in sympatry and allopatry when using different thresholds to define young species in lepidopterans and *Bufo* frogs. NC: regression did not converged and the analysis was not possible. Figure 3 shows similar results for premating and postzygotic isolation in *Drosophila.*

| Group | Cutoff | $c_{symp}$ | $c_{allop}$ | *W* | *P*-value |
|---|---|---|---|---|---|
| **Lepidopterans** | 0.5 | NC | NC | — | — |
| **Lepidopterans** | 1 | NC | NC | — | — |
| ***Bufo* frogs** | 0.05 | 0.028 | >0.05 | 995,004 | <0.0001 |
| ***Bufo* frogs** | 0.06 | 0.028 | >0.06 | 946,053 | <0.0001 |
| ***Bufo* frogs** | 0.07 | 0.028 | 0.037 | 816,440 | <0.0001 |
| ***Bufo* frogs** | 0.08 | 0.029 | >0.08 | 971,028 | <0.0001 |
| ***Bufo* frogs** | 0.09 | 0.028 | 0.037 | 839,586 | <0.0001 |
| ***Bufo* frogs** | 0.10 | 0.029 | 0.038 | 868,986 | <0.0001 |
| ***Bufo* frogs** | 0.11 | 0.029 | 0.038 | 869,901 | <0.0001 |
| ***Bufo* frogs** | 0.12 | 0.028 | 0.038 | 871,175 | <0.0001 |
